# Corticolimbic Circuit Structure Moderates an Association Between Early Life Stress and Later Trait Anxiety

**DOI:** 10.1101/581926

**Authors:** M. Justin Kim, Madeline J. Farber, Annchen R. Knodt, Ahmad R. Hariri

## Abstract

Childhood adversity is associated with a wide range of negative behavioral and neurodevelopmental consequences. However, individuals vary substantially in their sensitivity to such adversity. Here, we examined how individual variability in structural features of the corticolimbic circuit, which plays a key role in emotional reactivity, moderates the association between childhood adversity and later trait anxiety in 798 young adult university students. Consistent with prior research, higher self-reported childhood adversity was significantly associated with higher self-reported trait anxiety. However, this association was attenuated in participants with higher microstructural integrity of the uncinate fasciculus *and* greater thickness of the orbitofrontal cortex. These structural properties of the corticolimbic circuit may capture a neural profile of relative resiliency to early life stress, especially against the negative effects of childhood adversity on later trait anxiety. More generally, our findings highlight the potential utility in the simultaneous consideration of qualitatively different brain structural measures in explaining complex behavioral associations in future research.

## Introduction

Developmental neuroscience reveals widespread effects off childhood adversity on brain development, with a focus on neural systems that are responsible for the experience and regulation of negative emotions (Tottenham, 2014). Human neuroimaging studies have reported associations between childhood adversity and alterations in the functional activity, intrinsic connectivity, and structure of the corticolimbic circuit (Burghy et al., 2012; Dannlowski et al., 2012; Gold et al., 2016; Hanson et al., 2010; Kelly et al., 2013; Malter Cohen et al., 2013). The corticolimbic circuit, particularly the dynamic interactions between the amygdala and prefrontal cortex, is central to the emergence and regulation of emotion and anxiety (Casey et al., 2011; Kim et al., 2011; Ochsner & Gross, 2005). As research on childhood adversity and adult internalizing symptoms converges on the corticolimbic circuit recent studies have shifted their attention to elucidating neural mechanisms for the later emergence of anxiety or depression following childhood adversity (Casey et al., 2011; Gorka et al., 2014; Nusslock & Miller, 2016).

Two structural magnetic resonance imaging (MRI) measures have been frequently featured in this converging research: (a) the degree of structural connectivity between the amygdala and prefrontal cortex represented by the microstructural integrity of the uncinate fasciculus (UF), and (b) the thickness of the orbitofrontal cortex (OFC). In general, reduced OFC thickness and UF integrity are associated with increased internalizing symptoms and related behavioral/physiological characteristics, such as lesser fear extinction memory or greater trait anxiety (Eden et al., 2015; Greening & Mitchell, 2015; Hartley, 2011; Kim & Whalen, 2009; Kühn et al, 2011; Milad et al., 2005; but also see Ducharme et al., 2014). A dual processing framework is often used to explain these findings, such that limbic areas of the brain (e.g., amygdala) communicate with the prefrontal cortical regions (e.g., OFC) via direct and indirect reciprocal neural pathways (Aggleton et al., 2015; Ghashghaei et al., 2007). Dynamic communication along these pathways in turn supports patterns of emotional reactivity (Bishop et al., 2007; Kim et al., 2011; Ochsner & Gross, 2005; Quirk & Beer, 2006). As such, abnormalities of the corticolimbic circuit lead to an imbalance between the amygdala and the prefrontal cortex, which is proposed to be an underlying neural mechanism for exaggerated internalizing symptoms seen in affective disorders (Grupe & Nitschke, 2013; Helm et al., 2018; Shin & Liberzon, 2010). Indeed, studies suggest perturbations of the corticolimbic circuit may be a hallmark of many clinical disorders, including social anxiety disorder, generalized anxiety disorder, posttraumatic stress disorder, and major depressive disorder (Johnstone et al., 2007; LeWinn et al., 2014; Phan et al., 2009; Sadeh et al., 2016; Tromp et al., 2012).

Thus far, the majority of studies in this area have not considered the link between childhood adversity and anxiety later in life through the concurrent utilization of both structural connectivity and cortical thickness measures. While it is generally suggested that stronger corticolimbic structural connectivity and thicker OFC is associated with reduced internalizing symptoms, the *interactions* between these neural measures are seldom investigated. Examining such interactions is important because they allow for approximating the extent to which the capacity for both local regulatory processing via the OFC and dynamic communication between the OFC and amygdala shape emotional reactivity. The present study aims to address this need and identify a neural profile of those most resilient to the detrimental effects of childhood adversity on trait anxiety. Such a profile could subsequently inform ongoing efforts to develop biomarkers of relative risk or resilience for psychopathology as well as targets of intervention. Using data from 798 young adult university students through the Duke Neurogenetics Study, we specifically tested the hypothesis that individuals with higher microstructural integrity of the UF *and* thicker OFC would exhibit an attenuated link between childhood adversity and trait anxiety in adulthood.

## Methods

### Participants

Data were available from 798 undergraduate students (452 women, age range 18-22 years, mean age = 19.65 years) who successfully completed the Duke Neurogenetics Study (DNS) between January 25^th^, 2010 and November 12^th^, 2013 including structural and diffusion magnetic resonance imaging scans, as well as self-reported childhood stress and trait anxiety. All participants provided written informed consent according to the Duke University Medical Center Institutional Review Board. To be eligible for the DNS, participants were required to be free of the following conditions: 1) medical diagnoses of cancer, stroke, head injury with loss of consciousness, untreated migraine headaches, diabetes requiring insulin treatment, chronic kidney, or liver disease; 2) use of psychotropic, glucocorticoid, or hypolipidemic medication; and 3) conditions affecting cerebral blood flow and metabolism (e.g., hypertension).

As the DNS seeks to examine broad variability in multiple behavioral phenotypes related to psychopathology, participants were not excluded based on diagnosis of past or current Diagnostic and Statistical Manual of Mental Disorders, Fourth Edition (DSM-IV; American Psychiatric Association, 1994) Axis I or select Axis II (borderline and antisocial personality) disorder. However, all participants were not taking psychotropic medications for a minimum of 14 days before study initiation. Categorical diagnosis was assessed with the electronic Mini International Neuropsychiatric Interview (Lecrubier et al., 1997) and Structured Clinical Interview for the DSM-IV subtests (First et al., 1996). Of the total sample reported here, 153 participants (19%) met criteria for at least one current or past DSM-IV diagnosis. Detailed summary of diagnoses is presented in Supplemental Table S1 in the Supplemental Materials.

### Self-Report Questionnaires

The State-Trait Anxiety Inventory-Trait Version (STAI-T) was used to assess self-reported levels of trait anxiety (Spielberger et al., 1988). The Childhood Trauma Questionnaire (CTQ) was used to assess exposure to childhood stress in five categories: emotional abuse, physical abuse, sexual abuse, emotional neglect, and physical neglect (Bernstein et al., 1997). CTQ items were summed to create a total score for childhood adversity.

### Image Acquisition

Each participant was scanned using one of the two identical research-dedicated GE MR750 3T scanner equipped with high-power high-duty-cycle 50-mT/m gradients at 200 T/m/s slew rate, and an eight-channel head coil for parallel imaging at high bandwidth up to 1 MHz at the Duke-UNC Brain Imaging and Analysis Center. Following an ASSET calibration scan, diffusion-weighted images were acquired across two consecutive 2-min 50-s providing full brain coverage with 2 mm isotropic resolution and 15 diffusion-weighted directions (echo time (TE) = 84.9 ms, repetition time (TR) = 10,000 ms, *b* value = 1,000 s/mm^2^, field of view (FOV) = 240 mm, flip angle = 90°, matrix = 128 × 128, slice thickness = 2 mm. High-resolution anatomical T1-weighted MRI data were obtained using a 3D Ax FSPGR BRAVO sequence (TE = 3.22 ms, TR = 8.148 ms, FOV = 240 mm, flip angle = 12°, 162 sagittal slices, matrix =256 × 256, slice thickness = 1 mm with no gap).

### Diffusion MRI Analysis

All diffusion MRI data were preprocessed in accordance with the protocol developed by the Enhancing Neuro Imaging Genetics through Meta-Analysis consortium (ENIGMA; http://enigma.ini.usc.edu/protocols/dti-protocols). As these are standardized procedures made available by the ENIGMA consortium and further documented in our previously published work (Kim et al., 2019), a full description of diffusion MRI data preprocessing steps are provided in the Supplemental Methods available online. Following preprocessing, *a priori* pathways-of-interest were taken from the Johns Hopkins University DTI-based white matter atlas, adhering to the ENIGMA protocol (Wakana et al., 2007). These pathways-of-interest were used to mask each participant’s FA skeleton maps, and then the average FA values were extracted on an individual participant basis. Our primary pathway-of-interest was the uncinate fasciculus (Figure 1A). As the UF included in this atlas only represents a very small intermediary segment of the pathway, we used another UF pathway-of-interest with better coverage of the entire tract using the Johns Hopkins University White Matter Tractography Atlas (Mori et al., 2005). The left and right UF pathways were binarized in order to extract mean FA values for each participant. Since we did not have *a priori* predictions regarding inter-hemispheric differences and to reduce the number of statistical tests, the extracted FA values were averaged across left and right hemispheres for further statistical analyses to protect against false positives.

**Figure 1.**
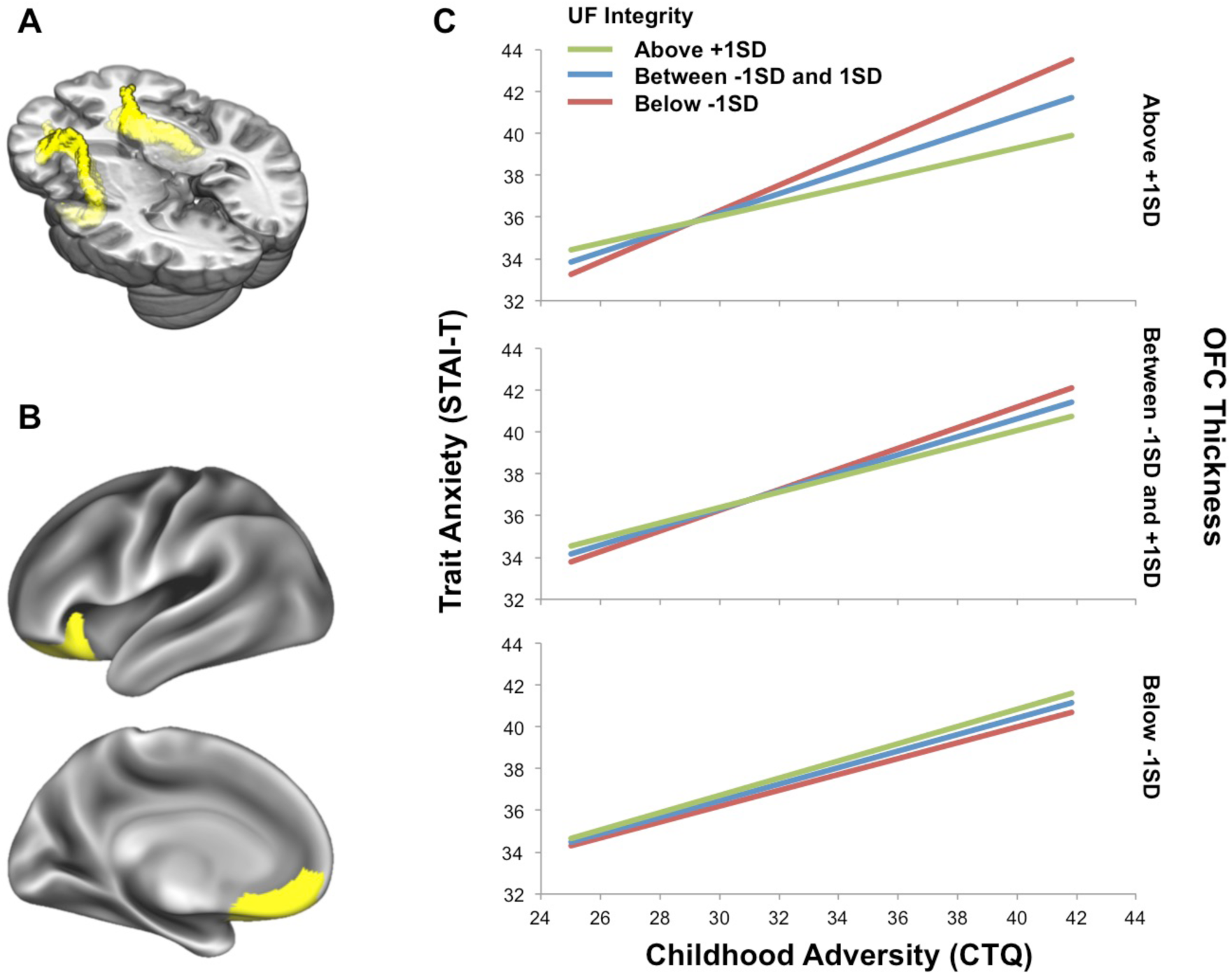
Corticolimbic circuit structure moderates the association between childhood adversity and trait anxiety in adulthood. (A) The uncinate fasciculus (UF), a major white matter pathway that connects corticolimbic circuit nodes, notably the amygdala and OFC, is depicted in yellow. Mean fractional anisotropy (FA) values were used to estimate the microstructural integrity of the uncinate fasciculus. (B) The orbitofrontal cortex is overlaid on a medial and lateral surface of an inflated brain image. Mean cortical thickness measures were estimated from these regions. (C) A significant three-way interaction was observed such that the microstructural integrity of the uncinate fasciculus and orbitofrontal cortical thickness jointly moderated the association between childhood adversity and adult trait anger. Attenuation of a usually robust association between childhood adversity and trait anxiety (bottom and middle panel) was observed in individuals with stronger microstructural integrity of the uncinate fasciculus *and* thicker orbitofrontal cortex (i.e., decreased slope of the green lines in the top panel).

### Cortical Thickness Analysis

To generate cortical thickness measures, T1-weighed images from all participants were first skull-stripped using ANTs (Klein et al., 2009), then submitted to FreeSurfer v.5.3 (http://surfer.nmr.mgh.harvard.edu)’s automated surface-based pipeline, *recon-all* (Dale et al., 1999; Fischl et al., 1999). Each brain volume’s surface underwent reconstruction and cortical parcellation (Klein & Tourville, 2012). From the reconstructed surface, cortical thickness was estimated from the distance between the gray/white matter boundary and the gray matter/cerebral spinal fluid boundary at each vertex (Fischl & Dale, 2000). Based on the *a priori* hypothesis, our primary region-of-interest was the orbitofrontal cortex. As the Desikan-Killany-Tourville atlas (Klein & Tourville, 2012) provides medial and lateral OFC (mOFC and lOFC, respectively) parcellations separately, cortical thickness of the OFC was computed as the average of the left and right mOFC and lOFC (Figure 1B). *Post hoc* analyses were performed using cortical thickness measures derived from the mOFC and lOFC separately.

### Statistical Analysis

PROCESS for SPSS (Hayes, 2013) was utilized within SPSS 21 (IBM Corp., Armonk, NY, USA) to test whether an interaction between microstructural integrity of UF and cortical thickness of the OFC jointly moderated the association between CTQ (independent variable) and STAI-T scores (dependent variables). Age, sex, and head movement during the diffusion MRI scans were included in the model as covariates. The latter was included as a covariate as it has been suggested that head motion may induce spurious correlations between diffusivity- or anisotropy-related metrics with behavioral measures (Yendiki et al., 2013). Current or past diagnosis of DSM-IV disorders was also included as a covariate, due to our interests in assessing the interactions among related variables that include trait anxiety and corticolimbic structures. In addition to the UF, the cingulum bundle was used as a control pathway-of-interest to test the moderation model with the same parameters to assess the possibility that the observed effects may be due to the variance associated with limbic white matter tracts in general. In a similar manner, the rostral anterior cingulate cortex was used as a control region-of-interest to examine the possibility that the observed effects may be due to limbic cortical thickness in general. Furthermore, amygdala volume, derived from FreeSurfer’s automated segmentation pipeline, was entered as one of the moderators in the model to test the possibility that the results are driven by cortical and subcortical structural variation. Finally, all of the analyses detailed above were repeated in participants without current or past DSM-IV diagnoses (*n* = 645).

## Results

### Self-Report Measures of Childhood Adversity and Trait Anxiety

Descriptive statistics (mean ± standard deviation) of the self-report measures are summarized as follows: CTQ (33.42 ± 8.4) and STAI-T (37.84 ± 9.19). Mean scores for the CTQ subscales were as follows: emotional abuse (7.13 ± 2.73), physical abuse (6.06 ± 1.98), sexual abuse (5.28 ± 1.49), emotional neglect (8.44 ± 3.55), and physical neglect (6.51 ± 2.24). As expected, a significant positive correlation was observed between CTQ and STAI-T (*r* = 0.42, *p* < 0.00001). This association remained after controlling for age, sex, and diagnosis of DSM-IV disorders, using hierarchical regression analysis with CTQ as an independent variable that was added in in the second step and STAI-T as the dependent variable (first step: *R*^2^ = 0.037, *F*(3,794) = 10.28, *p* < 0.000001; second step: **Δ***R*^2^ = 0.162, **Δ***F*(1,793) = 160.71, *β* = 0.41, *p* < 0.000001).

### Moderation Analysis

The overall model was significant in predicting trait anxiety from childhood adversity, microstructural integrity of the UF, and cortical thickness of the OFC (*R*^2^ = 0.21, *F*_(11, 786)_ = 19.23, *p* < 0.0001). Importantly, a significant three-way interaction among CTQ total scores, UF FA, and OFC cortical thickness predicted STAI-T scores (*b* = −26.58, 95% confidence interval (CI) = [−47.71, −5.45], **Δ***R*^2^= 0.006, *p* = 0.014). Follow-up simple slopes analysis showed that the interaction was primarily driven by individuals with relatively high UF FA and thicker OFC, for whom the association between CTQ and STAI-T scores was attenuated (*b* = 0.32, CI = [0.18, 0.47], *p* < 0.0001; Figure 1C). Specifically, the Johnson-Neyman technique indicated that the interaction between CTQ total scores and UF FA was significantly associated with STAI-T only when OFC cortical thickness was 0.09 standard deviations above the mean or greater. Conditional effects of CTQ on STAI-T at below −1 SD, between −1 SD and +1 SD, and above +1 SD of UF FA and OFC cortical thickness are summarized in Supplemental Table S2 in the Supplemental Material available online. This interaction was robust to the inclusion of global FA (i.e., grand mean FA of all voxels in the whole brain) as an additional covariate in the model (*b* = − 26.63, CI = [−47.78, −5.48], **Δ***R*^2^= 0.006, *p* = 0.014). When the cingulum bundle was used as a moderator variable instead of the UF, the three-way interaction was no longer statistically significant, regardless of the inclusion of the global FA covariate. Similarly, when the rostral anterior cingulate cortex thickness was used as a moderator variable instead of OFC, the three-way interaction was no longer statistically significant, regardless of the inclusion of the global FA covariate. When amygdala volume was entered into the models as one of the moderators (replacing UF FA or OFC thickness, respectively) with intracranial volume as an additional covariate, the three-way interactions were no longer significant. Finally, when moderation models were generated using only one of the two proposed moderators, the interaction was no longer significant, indicating that both UF FA and OFC thickness were necessary to observe a moderating effect.

Similar findings were observed when cortical thickness was estimated separately for the mOFC and lOFC. In brief, significant three-way interactions among CTQ total scores, UF FA, and medial/lateral OFC cortical thickness predicted STAI-T scores (model using mOFC: *b* = −21, CI = [−41.28, −0.72], **Δ***R*^2^= 0.004, *p* = 0.043; model using lOFC: *b* = −23.91, CI = [−42.38, −5.44], **Δ***R*^2^= 0.007, *p* = 0.011). In both cases, follow-up simple slopes analysis showed that the interaction was primarily driven by individuals with relatively high UF FA and thicker OFC (model using mOFC: *b* = 0.35, CI = [0.2, 0.51], *p* < 0.0001; model using lOFC: *b* = 0.31, CI = [0.16, 0.45], *p* < 0.0001). The Johnson-Neyman technique indicated that the interactions between CTQ total scores and UF FA were significantly associated with STAI-T, only when mOFC thickness was 0.34 standard deviations above the mean or greater, or when lOFC cortical thickness was 0.1 standard deviations above the mean or greater. All of the results remained consistent when the analyses were limited to 645 individuals without current or past DSM-IV diagnoses. Detailed description of these findings is summarized in the Supplemental Results.

## Discussion

Our findings offer further evidence that the link between the experience of childhood adversity and later trait anxiety is influenced by structural characteristics of the corticolimbic circuit. By leveraging two distinct structural properties of the corticolimbic circuit in the form of microstructural integrity of the UF and cortical thickness of the OFC, we observed a three-way interaction among these two measures and childhood adversity in predicting trait anxiety. Unpacking this interaction revealed that, while the association between childhood adversity and trait anxiety was generally robust, it was relatively attenuated in individuals who had both stronger microstructural integrity of the UF and thicker OFC.

These patterns inform neuroscience research that seeks to elucidate the developmental pathways through which the experience of childhood adversity may later emerge as differences in mood and affect. Cortical thinning of the OFC has been associated with maltreatment in children (Kelly et al., 2013), and reduced OFC thickness has been reported in adolescents who experienced childhood abuse (Gold et al., 2017). Relatedly, reduced volume of the OFC measured during adulthood has been associated with childhood adversity (Clausen et al., 2019; Dannlowski et al., 2012). Consistent with these individual findings, a voxel-wise meta-analysis of data from 693 children, adolescents, and adults found evidence of reduced OFC volume as a function of childhood maltreatment (Lim et al., 2014). A common implication from these findings is that structural properties of the OFC – and more generally, the corticolimbic circuit – could represent biomarkers of vulnerability to stress and, conversely, a neural architecture for resilience. The potential for OFC thickness specifically as a biomarker is further supported by the observation that temperamental characteristics related to behavioral inhibition measured as early as 4 months of age are associated with thinner OFC in young adulthood (Schwartz et al., 2010).

Our findings expand upon this proposal by showing how a generally robust association between childhood adversity and trait anxiety is moderated by the cortical thickness of the OFC, along with the microstructural integrity of the UF – another promising structural risk biomarker (Eden et al., 2015; Greening & Mitchell, 2015; Kim & Whalen, 2009; see Mincic, 2015 for review). The present findings extend this existing literature by demonstrating that OFC thickness and UF integrity may need to be considered simultaneously to realize the value of corticolimbic circuit structure as a biomarker of relative risk or resiliency to the negative effects of childhood adversity. A plausible explanation of this buffering account is that the link between childhood adversity and trait anxiety is attenuated by a fully developed neural platform for better regulation of negative emotions (Casey et al., 2011; Ochsner & Gross, 2005).

Considering the suggested protective role of stronger corticolimbic structural connectivity, it is noteworthy that the three-way interaction may also have been influenced by individuals with weaker UF integrity and thicker OFC. We observed an increased association between childhood adversity and trait anxiety in individuals with weaker microstructural integrity of the UF and thicker OFC (red line in the top panel of Figure 1C and **Supplemental Figure S1**). Individuals with such patterns of corticolimbic structure contributed to the observed effects by showing a steeper slope between childhood adversity and trait anxiety. One possible interpretation is that corticolimbic structural connectivity may serve a role as a defensive mechanism against childhood adversity, which is activated only when the OFC has not sufficiently matured. The latter speculation is inferred from evidence for age-related monotonic decline in cortical thickness across the brain (Walhovd et al., 2017). This particular prediction would benefit from future studies employing a longitudinal approach.

Our study, of course, is not without limitations. First, due to a cross-sectional design, childhood adversity was measured retrospectively and relied on self-report. Thus, participants may have been subject to biased recall of their childhood experiences and their self-report influenced by current mood. This line of research can greatly benefit from future studies with a longitudinal design, such as sampling prospective measures of childhood adversity from multiple sources. Second, as our data were sampled from high functioning young adult university students, the generalizability of the findings reported here remains to be determined, especially with regard to those who have endured severe early life stress (e.g., institutionalization, abuse). Third, it should be noted that corticolimbic white matter pathways, including the UF, are amongst the slowest to mature, continuing to develop until well into the third decade of life (Lebel et al., 2008). Since the present sample consisted entirely of young adults between 18 and 22 years of age, the structural properties of the UF may still be changing. As such, our findings should be interpreted within such constraints in mind. Fourth, the STAI-T, despite its namesake, has met with some criticism that it may reflect the tendency to experience both anxiety and depression (Watson et al., 1995). Thus, when interpreting STAI-T, it may be useful to consider it as representing a general negative emotion (Grupe & Nitschke, 2013). Lastly, independent replication is necessary if these features of corticolimbic circuit structure are to be further considered as risk biomarkers. This is particularly important with regard to the unexpected increased association between childhood adversity and trait anxiety in those with thicker OFC and lower UF integrity.

These limitations notwithstanding, our current findings build upon and extend the literature on identifying neuroimaging-derived moderators of potentially long-term, detrimental effects of childhood adversity. Our work further offers novel evidence that a specific pattern of corticolimbic circuit structural properties may capture a neural profile of resiliency to the negative impact of childhood adversity on trait anxiety. More generally, our findings demonstrate that simultaneously considering qualitatively distinct structural measures, such as microstructural integrity of white matter pathways and thickness of cortical gray matter, may be useful in elucidating the nature of known associations among experiential and behavioral phenomena.

## Acknowledgements

We thank the Duke Neurogenetics Study participants as well as the staff of the Laboratory of NeuroGenetics. The Duke Neurogenetics Study was supported by Duke University and National Institutes of Health (NIH) grants R01DA031579 and R01DA033369. ARH is further supported by NIH grant R01AG049789. The Duke Brain Imaging and Analysis Center’s computing cluster, upon which all DNS analyses heavily rely, was supported by the Office of the Director, NIH under Award Number S10OD021480.

