## Supplemental Materials for "Corticolimbic Circuit Structure Moderates an Association Between Early Life Stress and Later Trait Anxiety"

### Supplemental Methods

*Image Acquisition*

Each participant was scanned using one of the two identical research-dedicated GE MR750 3T scanner equipped with high-power high-duty-cycle 50-mT/m gradients at 200 T/m/s slew rate, and an eight-channel head coil for parallel imaging at high bandwidth up to 1 MHz at the Duke-UNC Brain Imaging and Analysis Center. For dMRI data, following an ASSET calibration scan, diffusion-weighted images were acquired across two consecutive 2-min 50-s providing full brain coverage with 2 mm isotropic resolution and 15 diffusion-weighted directions (echo time (TE) = 84.9 ms, repetition time (TR) = 10,000 ms, $b$ value = 1,000 s/mm$^2$, field of view (FOV) = 240 mm, flip angle = 90°, matrix = 128 × 128, slice thickness = 2 mm). For fMRI data, first a semi-automated high-order shimming program was used to ensure global field homogeneity. A series of 34 interleaved axial functional slices aligned with the anterior commissure-posterior commissure plane were acquired for full-brain coverage using an inverse-spiral pulse sequence to reduce susceptibility artifacts (TR/TE/flip angle=2000 ms/30 ms/60; FOV=240mm; 3.75×3.75×4mm voxels; interslice skip=0). Four initial radiofrequency excitations were performed (and discarded) to achieve steady-state equilibrium. To allow for spatial registration of each participant's data to a standard coordinate system, high-resolution anatomical T1-weighted MRI data were obtained using a 3D Ax FSPGR BRAVO sequence (TE = 3.22 ms, TR = 8.148 ms, FOV = 240 mm, flip angle = 12°, 162 sagittal slices, matrix =256 × 256, slice thickness = 1 mm with no gap).

*dMRI Data Preprocessing*

All dMRI data were preprocessed in accordance with the protocol developed by the Enhancing Neuro Imaging Genetics through Meta-Analysis consortium (ENIGMA; http://enigma.ini.usc.edu/protocols/dti-protocols/). Raw diffusion-weighted images were corrected for eddy current and aligned to the non-diffusion-weighted (b0) image using linear registration in order to correct for head motion. Volume-by-volume head motion was quantified by calculating the root mean square (RMS) deviation of the six motion parameters (three translation and three rotation components), for each pair of consecutive diffusion-weighted brain volumes. The resulting volume-by-volume RMS deviation values were averaged across all images, yielding a summary statistic of head motion for each participant, which were used as a covariate of no interest in all subsequent group level analyses. Next, following skull stripping, diffusion tensor models were fit at each voxel using the Diffusion Toolbox in FSL (Behrens et al., 2003; Smith et al., 2004), generating a whole-brain fractional anisotropy (FA) image for each participant. For each participant, an average of the FA images from each of the two scans were produced to increase signal-to-noise ratio of the data. These images were then subjected to a tract-based spatial statistics (TBSS) in FSL (Smith et al., 2006). TBSS analysis entails the realignment of each individual FA image to a standard FA template in Montreal Neurological Institute (MNI) space using nonlinear registration (FNIRT). Then, individual

FA images were projected onto the ENIGMA-DTI FA skeleton, searching for maximal FA values perpendicular to the skeleton. The resulting individual FA skeletons were used to extract average FA values for our *a priori* pathways of interest.

### Supplemental Results

*Moderation Analysis in the Normative Sample (n = 645)*

Results remained consistent when the analyses were limited to 645 individuals without current or past DSM-IV diagnoses. The overall model was significant in predicting trait anxiety from childhood adversity, UF integrity, and OFC thickness ($R^2$ = 0.15, $F_{(10, 634)}$ = 10.93, $p$ < 0.0001). A significant three-way interaction among CTQ total scores, UF FA, and OFC thickness predicted STAI-T scores ($b$ = -29.85, CI = [-56.93, -2.78], $\Delta R^2$ = 0.006, $p$ = 0.031). Follow-up simple slopes analysis showed that the interaction was primarily driven by individuals with relatively high UF FA and thicker OFC, for whom the association between CTQ and STAI-T scores was attenuated ($b$ = 0.22, CI = [0.03, 0.4], $p$ = 0.022; **Supplemental Figure S1**). The Johnson-Neyman technique indicated that the interaction between CTQ total scores and UF FA was significantly associated with STAI-T only when OFC cortical thickness was 0.15 standard deviations below the mean or greater. Conditional effects of CTQ on STAI-T at below -1 SD, between -1 SD and +1 SD, and above +1 SD of UF FA and OFC cortical thickness are summarized in Table S3 in the Supplemental Material available online. This interaction was robust to the inclusion of global FA (i.e., grand mean FA of all voxels in the whole brain) as an additional covariate in the model ($b$ = -29.17, CI = [-56.25, -2.09], $\Delta R^2$ = 0.006, $p$ = 0.035). All control analyses were consistent with the findings from the full sample – the inclusion of cingulum bundle, rostral anterior cingulate thickness, amygdala volume didn't yield significant results. No significant moderating effects from single moderator models were observed.

Once again, results remained consistent when cortical thickness was estimated separately for the mOFC and lOFC. Significant three-way interactions among CTQ total scores, UF FA, and medial/lateral OFC cortical thickness predicted STAI-T scores (model using mOFC: $b$ = -27.27, CI = [-53.88, -0.66], $\Delta R^2$ = 0.006, $p$ = 0.045; model using lOFC: $b$ = -23.61, CI = [-46.22, -5.44], $\Delta R^2$ = 0.006, $p$ = 0.041). In both cases, follow-up simple slopes analysis showed that the interaction was primarily driven by individuals with relatively high UF FA and thicker OFC (model using mOFC: $b$ = 0.24, CI = [0.05, 0.43], $p$ = 0.012; model using lOFC: $b$ = 0.2, CI = [0.01, 0.38], $p$ = 0.012). The Johnson-Neyman technique indicated that the interactions between CTQ total scores and UF FA were significantly associated with STAI-T, only when mOFC thickness was 0.07 standard deviations below the mean or greater, or when lOFC cortical thickness was 0.12 standard deviations below the mean or greater.

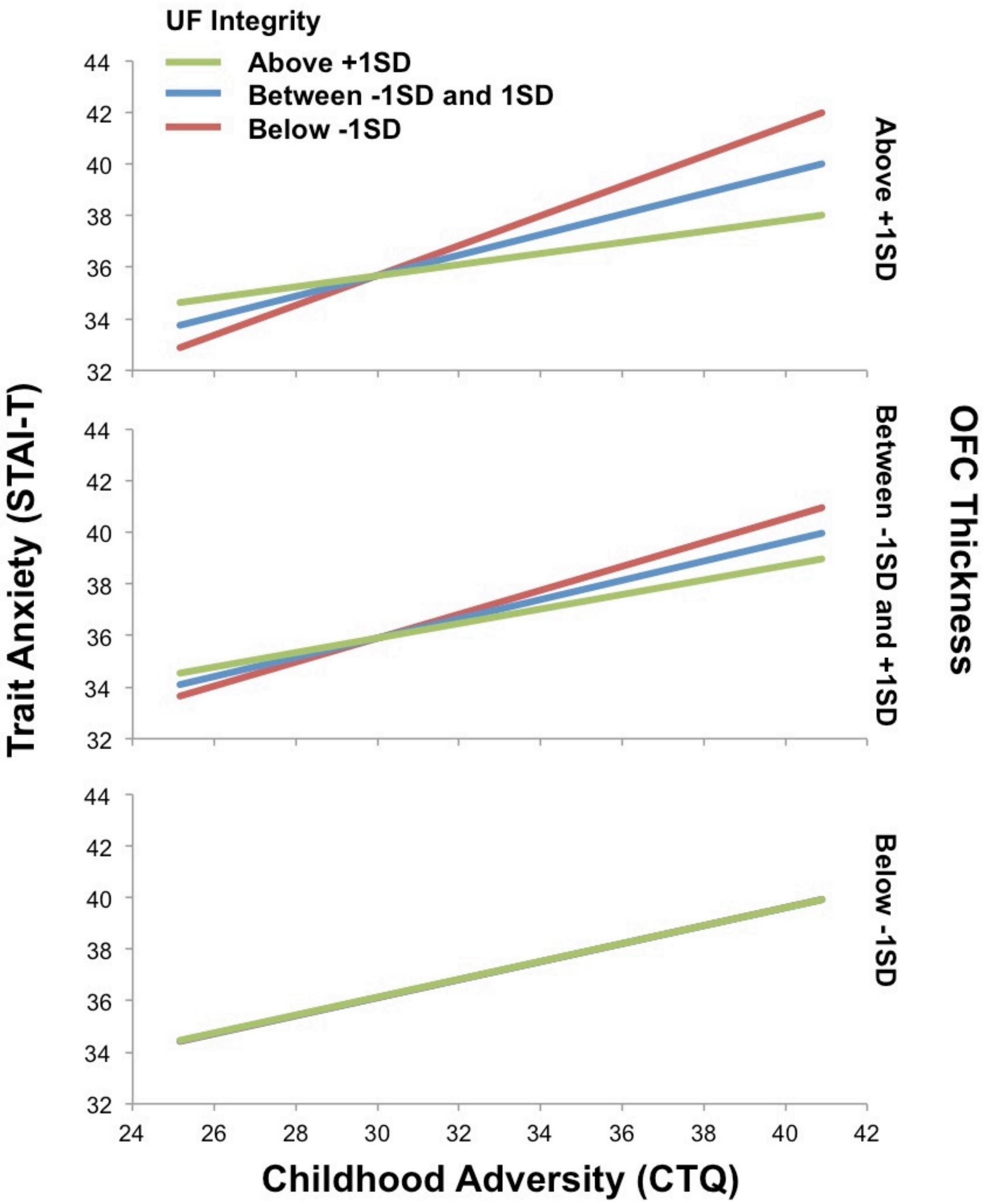


**Supplemental Figure S1.** Similar moderating effects were observed when the analysis was restricted to individuals with no current or past diagnoses of mental disorders *(n* =645). Attenuation of a usually robust association between childhood adversity and trait anxiety (bottom and middle panel) was observed in individuals with stronger microstructural integrity of the uncinate fasciculus *and* thicker orbitofrontal cortex (i.e., decreased slope of the green lines in the top panel).

**Supplemental Table S1.** DSM-IV diagnoses for current or past mental disorders in the total study sample (*n* = 153).

| Diagnosis | Count |
|---|---|
| Major depressive disorder | 25 |
| Bipolar disorder I or II | 5 |
| Bipolar disorder – Not otherwise specified | 10 |
| Hypomanic episode | 14 |
| Panic disorder | 9 |
| Agoraphobia | 13 |
| Social anxiety disorder | 7 |
| Generalized anxiety disorder | 16 |
| Obsessive-compulsive disorder | 7 |
| Posttraumatic stress disorder | 1 |
| Alcohol abuse/dependence | 93 |
| Substance (non-alcohol) abuse/dependence | 27 |
| Eating disorder | 6 |
| Antisocial personality disorder | 1 |
| Borderline personality disorder | 2 |

*Note*: The sum of the individual diagnoses is higher than the number of participants with diagnosed disorders because of multiple comorbid diagnoses. DSM-IV: Diagnostic and Statistical Manual of Mental Disorders, Fourth Edition.

**Supplemental Table S2.** Conditional effects of childhood adversity on trait anxiety at low, intermediate, and high levels of the UF integrity and OFC thickness (full sample; *n* = 798).

| OFC Thickness | UF Integrity | *b* (SE) | *t* | *p* |
|---|---|---|---|---|
| Low | Low | 0.38 (0.07) | 5.59 | < 0.0001 |
| Low | Intermediate | 0.40 (0.05) | 8.31 | < 0.0001 |
| Low | High | 0.41 (0.05) | 7.59 | < 0.0001 |
| Intermediate | Low | 0.49 (0.05) | 10.27 | < 0.0001 |
| Intermediate | Intermediate | 0.43 (0.04) | 12.20 | < 0.0001 |
| Intermediate | High | 0.37 (0.05) | 7.28 | < 0.0001 |
| High | Low | 0.61 (0.07) | 8.49 | < 0.0001 |
| High | Intermediate | 0.47 (0.05) | 9.34 | < 0.0001 |
| High | High | 0.32 (0.08) | 4.29 | < 0.0001 |

*Note*: Low, intermediate, and high was defined as < -1 standard deviation (SD), -1 SD < and < +1 SD, and > +1 SD, respectively. OFC: orbitrofrontal cortex; SE: standard error; UF: uncinate fasciculus.

**Supplemental Table S3.** Conditional effects of childhood adversity on trait anxiety at low, intermediate, and high levels of the UF integrity and OFC thickness (normative sample, *n* = 645).

| OFC Thickness | UF Integrity | *b* (SE) | *t* | *p* |
|---|---|---|---|---|
| Low | Low | 0.35 (0.08) | 4.62 | < 0.0001 |
| Low | Intermediate | 0.35 (0.05) | 6.52 | < 0.0001 |
| Low | High | 0.35 (0.07) | 4.87 | < 0.0001 |
| Intermediate | Low | 0.46 (0.05) | 8.57 | < 0.0001 |
| Intermediate | Intermediate | 0.37 (0.04) | 8.99 | < 0.0001 |
| Intermediate | High | 0.28 (0.06) | 4.56 | < 0.0001 |
| High | Low | 0.58 (0.09) | 6.66 | < 0.0001 |
| High | Intermediate | 0.40 (0.06) | 6.31 | < 0.0001 |
| High | High | 0.22 (0.09) | 2.29 | 0.0221 |

*Note*: Low, intermediate, and high was defined as < -1 standard deviation (SD), -1 SD < and < +1 SD, and > +1 SD, respectively. OFC: orbitrofrontal cortex; SE: standard error; UF: uncinate fasciculus.
